# JAGGED1 Stimulates Cranial Neural Crest Cell Osteoblast Commitment Pathways and Bone Regeneration Independent of Canonical NOTCH Signaling

**DOI:** 10.1101/2020.06.24.169755

**Authors:** Archana Kamalakar, Jay M. McKinney, Daniel Salinas Duron, Angelica M. Amanso, Samir A. Ballestas, Hicham M. Drissi, Nick J. Willett, Pallavi Bhattaram, Andrés J. García, Levi B. Wood, Steven L. Goudy

## Abstract

Craniofacial bone loss is a complex clinical problem with limited regenerative solutions. Currently, BMP2 is used as a bone-regenerative therapy in adults, but in pediatric cases of bone loss, it is not FDA-approved due to concerns of life-threatening inflammation and cancer. Development of a bone-regenerative therapy for children will transform our ability to reduce the morbidity associated with current autologous bone grafting techniques. We discovered that JAGGED1 (JAG1) induces cranial neural crest (CNC) cell osteoblast commitment during craniofacial intramembranous ossification, suggesting that exogenous JAG1 delivery is a potential craniofacial bone-regenerative approach. In this study, we found that JAG1 delivery using synthetic hydrogels containing O9-1 cells, a CNC cell line, into critical-sized calvarial defects in C57BL/6 mice provided robust bone-regeneration. Since JAG1 signals through canonical (*Hes1/Hey1*) and non-canonical (JAK2) NOTCH pathways in CNC cells, we used RNAseq to analyze transcriptional pathways activated in CNC cells treated with JAG1±DAPT, a NOTCH-canonical pathway inhibitor. JAG1 upregulated expression of multiple NOTCH canonical pathway genes (*Hes1*), which were downregulated in the presence of DAPT. JAG1 also induced bone chemokines (*Cxcl1*), regulators of cytoskeletal organization and cell migration (*Rhou*), signaling targets (STAT5), promoters of early osteoblast cell proliferation (*Prl2c2, Smurf1* and *Esrra*), and, inhibitors of osteoclasts (*Id1*). In the presence of DAPT, expression levels of *Hes1* and *Cxcl1* were decreased, whereas, *Prl2c2, Smurf1, Esrra, Rhou* and *Id1* remain elevated, suggesting that JAG1 induces osteoblast proliferation through these non-canonical genes. Pathway analysis of JAG1+DAPT-treated CNC cells revealed significant upregulation of multiple non-canonical pathways, including the cell cycle, tubulin pathway, regulators of *Runx2* initiation and phosphorylation of STAT5 pathway. In total, our data show that JAG1 upregulates multiple pathways involved in osteogenesis, independent of the NOTCH canonical pathway. Moreover, our findings suggest that JAG1 delivery using a synthetic hydrogel, is a bone-regenerative approach with powerful translational potential.

## 1. Introduction

Alagille syndrome manifests itself in patients as cardiac, biliary, and bony phenotypes, including increased incidence of fractures and bone loss (1). These patients also display abnormal vertebral segmentation and maxillary bone hypoplasia (2). Loss of function mutations in *JAGGED1 (JAG1)* have been implicated in Alagille syndrome (2, 3), suggesting that promotion of JAG1 signaling may stimulate bone induction which may be harnessed in the development of novel bone-regenerative therapies.

JAG1 is a membrane bound ligand of the NOTCH receptors 1-4 and signals via canonical (*Hes1/Hey1*) (4-7) and non-canonical (AKT, JAK2, MAPK and p38) intracellular pathways (8-12). JAG1 is required for normal craniofacial bone development in humans during which cranial neural crest (CNC) cells arise from the neural placode and migrate into the first brachial arch to form the calvarium and maxilla (4, 13). CNC cells then undergo cell fate commitment to become osteoblasts and form bone via intramembranous ossification (14, 15). Our prior research demonstrated that JAG1 signaling is required for CNC cell osteoblast commitment and that conditional ablation of *Jag1* in the CNC cells of mice was associated with craniofacial bone loss, recapitulating the facial phenotype seen in Alagille Syndrome patients who have *JAG1* mutations (4). JAG1 signaling occurs through cell-to-cell interactions between the JAG1 ligand and NOTCH receptors 1-4 (16-18) and we previously reported that JAG1-dependent CNC cell osteoblast commitment occurs primarily through non-canonical JAG1-JAK2 signaling (8).

The role of NOTCH signaling during osteoblast commitment and bone formation is both time and context dependent (19, 20). Our group and others have demonstrated the role and requirement of NOTCH signaling during bone development, and multiple studies have shown the importance of NOTCH signaling in human bone diseases (4, 21). The NOTCH canonical pathway plays an integral regulatory role in bone development and remodeling of the skeleton (22, 23). NOTCH signaling has been reported to either stimulate or inhibit osteogenesis depending on cell type and micro-environment (21). In particular, canonical NOTCH signaling has been shown to suppress the differentiation of skeletal cells, whereas NOTCH1 overexpression increases osteoblast commitment and mineralization in mesenchymal stem cells (24). Moreover, activation of *JAG1* is associated with increased bone mineral density and loss of function mutations in *JAG1* are associated with bone loss in patients with Alagille syndrome (25-27). These data suggest that the canonical NOTCH pathway is associated with bone loss while JAG1-associated bone formation occurs through the non-canonical NOTCH pathway. Based on our prior findings that JAG1 induces CNC cell differentiation into osteoblasts via a non-canonical NOTCH pathway mediated by phosphorylated JAK2 (8), we sought to determine the canonical vs non-canonical roles of JAG1-NOTCH signaling in CNC cells to identify potential novel therapeutic candidates for bone regeneration.

Therapies involving delivery of proteins or cells alone to regenerate bone have not reliably demonstrated regenerative capabilities, except for BMP2 (28-31). For decades, allografts and autografts of bone have been used to facilitate bone regeneration, however these techniques are expensive and are associated with significant morbidity (32, 33). Allografts have a decreased regenerative capacity and can lead to disease transmission from donor to recipient (32). Autografts reduce the risk of disease transmission and provide improved regenerative capacity but have significant morbidity at the donor site (33). Biopolymer-based drug and cell delivery therapies have gained popularity for tissue regeneration (34). JAG1 is a cell surface transmembrane protein and must be immobilized within a delivery scaffold for NOTCH signaling to occur (27).

In this study we demonstrate the ability of JAG1 delivery using a synthetic hydrogel encapsulating CNC cells to induce bone regeneration in a critical-sized calvarial defect and identify a novel non-canonical signaling pathway downstream of JAG1-JAK2-STAT5 using RNAseq during CNC cell osteoblast commitment. Together our findings reveal that exogenous delivery of JAG1 is a highly novel approach to promote non-canonical NOTCH signaling, ossification pathways and craniofacial bone regeneration.

## 2. Material and Methods

### 2.1. Cell culture

O9-1 cells, a stable mouse CNC cell line, (35) (Millipore sigma, SCC049) were seeded on a layer of Matrigel (Fisher, CB-40234) (1:50 dilution in 1X PBS). The cells were maintained and passaged in T75 flasks, using embryonic stem cell medium (Sigma, ES-1010-b) + 1% antibiotics (Penicillin/Streptomycin) (Gibco, 15240-062).

### 2.2. JAG1 immobilization

50 μL (1.5 mg) of Dynabeads Protein G (36) (Invitrogen 10003D) were transferred to a tube, where the beads were separated from the solution using a magnet. Recombinant JAG1-Fc (5 µM, 5.7 µM, 10 µM or 20 µM) (Creative Biomart, JAG1-3138H) and IgG-Fc fragment (5 µM or 5.7 µM) (Abcam, ab90285) alone were diluted in 200 μL PBS with 0.1% Tween-20 (Fisher, BP337-500) and then added to the dynabeads. The beads plus proteins were incubated at 4°C with rotation for 16 hours. Thereafter, the tubes were placed on a magnetic rack and the supernatant was removed. The bead-Ab complex was resuspended in 200 μL PBS with 0.1% Tween-20 to wash by gentle pipetting. The wash buffer was also separated from the beads-Ab complex using the magnet and the final suspension of the beads in hydrogels was used as treatment.

### 2.3. Hydrogel preparation

Poly (ethylene glycol) (PEG)-based synthetic hydrogels incorporating cell adhesive peptides were formed in two steps. First, maleimide end-functionalized 20 kDa four-arm PEG macromer (PEG-4MAL, > 95% end-group substitution, Laysan Bio, 4ARM-PEG-MAL-20K), was reacted with a thiol-containing adhesive peptide GRGDSPC (Genscript, RP20283) in PBS with 4 mM TEA at pH 7.4 for 1 hour. Then, the RGD-functionalized PEG-4MAL macromers were cross-linked in the presence of CNC cells and JAG1-dynabeads into a hydrogel by addition of the dithiol protease-cleavable peptide cross-linker GPQ-W (GCRDGPQGIWGQDRCG) (New England Peptides, Inc, (NEP) Custom made) (34). The final gel formulation consisted of 4% wt/vol polymer and 2.0 mM RGD.

### 2.4. *In vivo* experiments

All *in vivo* experiments were performed using procedural guidelines with appropriate approvals from the Institutional Animal Care and Use committee of Emory University. 6-8-week-old male and female C57BL/6 mice (The Jackson Laboratory, 000664) were used. The mice were housed in groups of 3. As shown in **Fig. 1B-G**, incisions were made using sterile surgical equipment to expose the parietal bones of the mice. The exposed surgery site was disinfected and 3.5 mm defects were created in the parietal bones using a variable speed drill (Dremel 3000) and sterile drill bits. **JAG1 Delivery:** 20 µls of 4% wt/vol PEG-4MAL hydrogels incorporated with JAG1-dynabeads without (n = 3 for all treatment groups) or with 100,000 O9-1 cells (n = 5 for all treatment groups) were placed within the defects created in parietal bones in the C57BL/6 mouse skulls as the 1^st^ dose. The 2^nd^ and 3^rd^ doses of the hydrogels encapsulating CNC cells and dynabead-bound JAG1 were administered as transcutaneous injections during week 4 and week 8, respectively. 2 mice died after 1^st^ dose of BMP2 + CNC cells treatment and 3 after Dynabeads + CNC cells treatment. The skulls were then harvested at week 12 and subjected to micro computed tomography.

**Figure 1:**
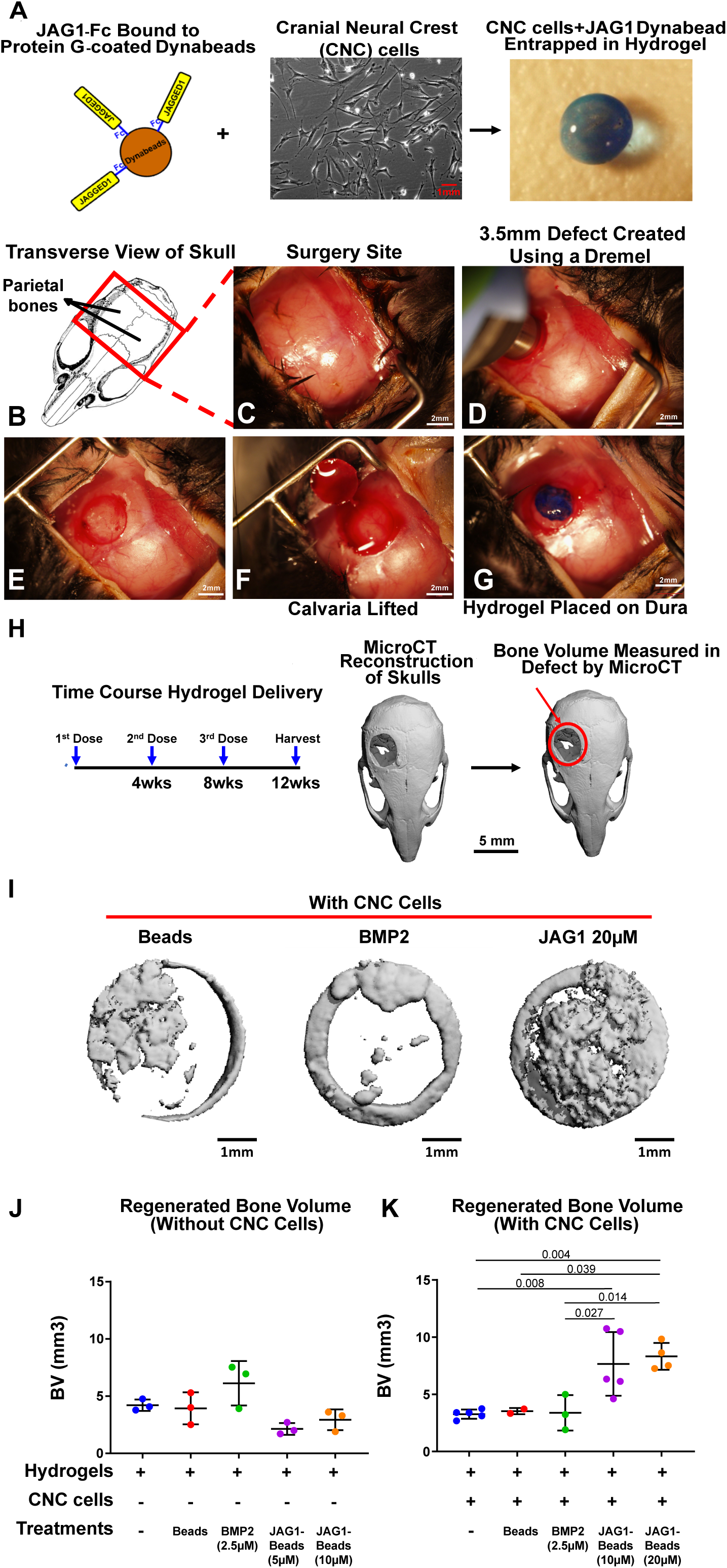
JAG1 delivery in a PEG hydrogel stimulates bone regeneration in a critical-sized bone defect mouse model. As a proof of concept experiment, we (A) incorporated JAG1-dynabeads complex (5 μM, 10 μM or 20 μM), dynabeads alone and BMP2 (2.5 µM) in 4% PEG-MAL hydrogels and implanted them into (B-G) 3.5 mm critical-sized defects in the parietal bones of 6-8-week old C57BL/6 mice and delivered them (J) without or (K) with CNC cells into the defects as 3 separate doses (Initial dose, Week 4, Week 8), as shown in H. After 12 weeks, we quantified differences in regenerated bone volume (J-K) within the defect and compared them between experimental groups by μCT analysis. μCT reconstructions of defects are shown in I. Data were subjected to ANOVA and Tukey’s post-test and are presented as mean (n ≥ 2) ± SD with p values reported.

### 2.5 Micro Computed Tomography (µCT)

µCT analyses were conducted according to the current guidelines for the assessment of bone volume within the defects created in mouse calvaria. Briefly, formalin-fixed skulls were positioned in the µCT tubes with nose facing the bottom of the tube and imaged using a µCT 40 (Scanco Medical AG, Bassersdorf, Switzerland) using a 36 µm isotropic voxel size in all dimensions. Thereafter, using a consistent and pre-determined threshold of 55 kVp, 145 µA, 8 W and 200 ms integration time for all measurements, three-dimensional (3D) reconstructions were created by stacking the regions of interest from ∼600 two-dimensional (2D) slices consisting of the entire skull and then applying a gray-scale threshold of 150 and Gaussian noise filter (σ = 0.8, support = 1.0), a coronal reformatting was done. Thereafter, a circular region of interest (ROI) encompassing the defect was selected for analysis consisting of transverse CT slices encompassing the entire defect and new bone volume (BV) was calculated.

### 2.6. RNA Sequencing

O9-1 cells were cultured in 24-well culture plates. When cells reached 100% confluency, they were treated (n = 3) for 12 hours with Fc-dynabeads (5.7 µM), unbound BMP2 (100 nM) (Creative biomart, BMP2-01H) and JAG1-dynabeads (5.7 µM) with or without DAPT (15 µM) (Millipore Sigma, D5942). After 12 hours, RNA was isolated using the Qiagen RNeasy kit (Qiagen, 74106) according to manufacturer’s protocols. The samples were submitted to the Molecular Evolution core at the Georgia Institute of Technology for sequencing. Quality Control (QC) was done using a bioanalyzer to determine that the RNA Integrity Number (RIN) of the samples was greater than 7. Thereafter a New England Biolab’s (NEB) mRNA isolation module in conjunction with their Ultra II RNA directional kit (NEB, E7760) was used to generate libraries for sequencing. QC was then performed on these libraries using the bioanalyzer and the library was quantified using fluorometric methods. Paired-end 75 base pairs (PE75) sequencing was performed on the Illumina NEXTSeq instrument to obtain a sequencing depth of 30 million reads per sample. The transcripts obtained were aligned using the mm9 genome reference database along with elimination of duplicate reads, using the strand next generation sequencing (NGS 3.4) application. The RNA levels were calculated in reads per kilobase per million mapped reads (RPKM). Genes expressed at > 1.5 RPKM were retained for further analyses (37).

### 2.7: Gene Set Variation Analysis

To establish differences in each gene set, we used gene set variation analysis (GSVA v1.36.1 package in R) to identify enrichment of each gene set across all samples (38). GSVA is an improved gene set enrichment method that detects subtle variations of pathway activity over a sample population in an unsupervised manner. The GSVA was conducted using the Molecular Signatures Database C2 gene sets v7.0 (MSigDB) (39). Statistical differences in enrichment scores for each gene set between groups were computed by comparing the true differences in means against the differences computed by a random distribution obtained by permuting the gene labels 1000 times. False discovery rate (FDR) adjusted q-values were computed using the method by Benjamini & Hochberg (40). Gene sets with FDR adjusted q-values < 0.05 were considered significant.

### 2.8. STAT multiplex array

O9-1 cells were cultured in 24-well culture plates. When cells reached 100% confluency, they were treated (n = 3) with Fc-dynabeads (5.7 µM), unbound BMP2 (100 nM) and JAG1-dynabeads (5.7 µM) with or without DAPT (15 µM) for 30 minutes and then lysed to obtain whole cell proteins (2 µg). The lysates were then probed for phosphorylated STATs using the Milliplex map STAT cell signaling magnetic bead 5-Plex kit (Millipore Sigma, 48-610MAG) according to manufacturer’s protocol.

### 2.9. Statistics

Bone data were analyzed by analysis of variance (ANOVA) with Tukey’s post-test using GraphPad Prism 8. All data are presented as mean (n ≥ 2 samples) ± SD. p < 0.05 between groups was considered significant and are reported as such.

### 2.10. Volcano plots

Differentially expressed genes were identified via volcano plot filtering. Volcano plots were constructed in R using the Enhanced Volcano package v. 1.6.0. Filtering threshold for expression difference presented as ‘log_2_ fold change’ was set at 1.5-fold change (log_2_ (1.5) = 0.585) and thresholding for significance, presented as ‘-log_10_ P’, was set at α=0.05 (-log_10_(0.05) = 1.301). For the comparison of JAG1 and JAG1 + DAPT treated samples, we assessed only the genes which were above the thresholds set for expression difference and significance value, (fold change ≥ 1.5 and p ≤ 0.05) when comparing these two respective groups to the Fc control.

### 2.11 PCA

Principal component analysis (PCA) was conducted in R using the stats package v.3.6.2. An orthogonal rotation in the Principal Component (PC) 1 – PC 2 plane was used to obtain new PC’s that better separated treatment groups.

## 3. Results

### JAG1 Stimulation of Cranial Neural Crest Cells in a PEG Hydrogel Repairs Cranial Defects

Delivery of unbound JAG1 or CNC cells alone to critically sized calvarial defects has not successfully regenerated bone due to low protein stability, cell survival, and decreased dosing kinetics compared to when delivered in synthetic materials (28). Thus, we sought to determine whether calvarial bone regeneration can occur with the sustained delivery of JAG1 with CNC cells. To do so, we developed a synthetic PEG hydrogel to encapsulate and deliver JAG1 and CNC cells. PEG hydrogels are made of four-arm PEG macromer that are end-functionalized by maleimide moieties (PEG-4MAL) to improve ligand binding and cross-linking. These hydrogels have been used extensively for targeted protein delivery because they are safe, elicit minimal inflammation, and have tunable properties (28). Because soluble JAG1 delivery has been found to inhibit, rather than stimulate NOTCH signaling (41), we immobilized a functional recombinant JAG1-IgG Fc fragment construct on to protein G-coated dynabeads (JAG1) with JAG1 exposed on the surface, as previously published (36). JAG1-dynabeads were then encapsulated within a 4% PEG-4MAL hydrogel together with CNC cells (**Fig. 1A**). To test the regenerative capacity of the JAG1-PEG-4MAL-CNC hydrogel, we implanted them in critical-sized parietal bone defects (3.5 mm) in mice (**Fig. 1B-G**). The volume of bone regenerated by the JAG1-PEG-4MAL-CNC hydrogel with 5 µM, 10 µM or 20 µM JAG1 was measured and compared to hydrogel alone, dynabeads alone, and BMP2 (2.5 µM) treatments, with or without CNC cells. To maintain continuous osteogenic induction, additional JAG1-PEG-4MAL-CNC hydrogels were injected into the defects transcutaneously at week 4, and week 8 (**Fig. 1H**). After 12 weeks, we quantified differences in bone volume (BV) within the cranial defect using μCT analysis. μCT reconstruction of defects demonstrated that in the absence of CNC cells, BMP2 was able to regenerate bone to reduce the critical size of the calvarial defects (**Fig. 1J & Supp. Fig 1**). The delivery of JAG1-PEG-4MAL-CNC hydrogels demonstrated significantly improved closure of the cranial defect at 20 µM JAG1 and 10 µM JAG1 had less but significant amount of bone regeneration too (**Fig. 1K**). Interestingly, the BMP2-PEG-4MAL-CNC hydrogels did not show increased bone regeneration (**Fig. 1K**). Together, these findings suggest that hydrogel delivery of JAG1 can induce CNC cell osteoblast commitment in order to repair cranial bone defects.

### JAG1 Up-regulates both Canonical and Non-Canonical Pathways in CNC cells

Previously, our lab demonstrated that JAG1 can elicit a NOTCH non-canonical JAK2 signal which stimulates CNC cell osteoblast commitment (8), however the downstream mediators are unknown. To determine the non-canonical downstream mediators of JAG1-JAK2 signaling, CNC cells (n = 3) were treated in the presence and absence of the canonical NOTCH inhibitor DAPT (15 µM) with JAG1 (5.7 µM) and compared to CNC cells treated with BMP2 (100 nM) and Fc (5.7 µM) as positive and negative controls, respectively. Cells were conditioned for 12 hours and then RNA was isolated. Gene expression was quantified for 10,016 genes using RNAseq (**Fig. 2A**).

**Figure 2:**
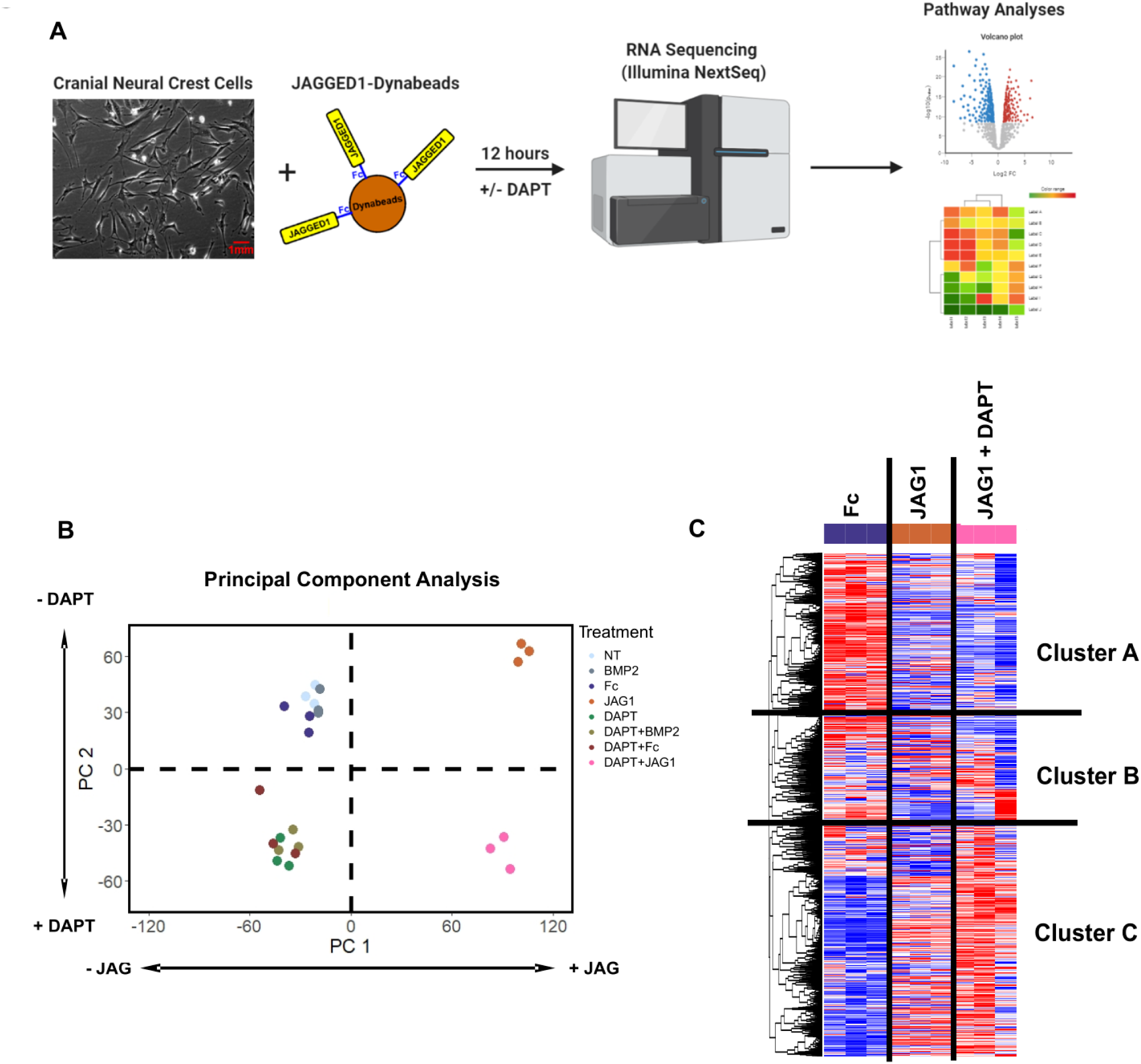
JAG1 induces canonical and non-canonical pathways in CNC cells. (A) CNC cells were treated with JAG1-dynabeads complex (5.7 μM), Fc-dynabeads complex (5.7 µM) and BMP2 (100 nM) with or without DAPT (15 µM), gene expression was quantified via RNAseq, and, univariate and pathway analysis was used to evaluate differences in gene expression. (B) A principal component analysis of all treatments revealed that PC1 distinguished JAG1-treated cells to the right from non-JAG1 treated cells to the left along the horizontal axis. PC2 distinguished cells treated with JAG1 alone towards the top and JAG1 + DAPT treated cells towards the bottom along the vertical axis. (C) Clustering analysis using Pearson correlation-coefficient reveals the heatmap showing distinct gene clusters downstream of different treatments (n = 3). Cluster A and B consists of genes downregulated by JAG1 and Cluster C consists of genes upregulated by JAG1.

To determine whether there were differences among the 8 treatment groups, we used a principal component analysis (PCA) to reduce the dimensionality of our dataset into two principal components (PCs, **Fig. 2B**). The analysis revealed that PC1 distinguished JAG1-treated cells to the right from non-JAG1 treated cells to the left along the horizontal axis. PC2 distinguished cells treated with JAG1 alone towards the top and cells co-treated with JAG1 and DAPT towards the bottom along the vertical axis. Since DAPT inhibits canonical, but not non-canonical NOTCH signaling, these findings indicate that JAG1 has a profound effect on CNC cell gene expression that is independent of canonical NOTCH signaling. Interestingly, these data also show that BMP2-treated cells grouped together with no-treatment (NT) and Fc controls, suggesting, as shown before, that although BMP2 signaling is essential for the formation and migration of CNC cells, BMP2 does not induce these precursors towards intramembranous ossification (42).

To determine the downstream targets of non-canonical JAG1 signaling, we compared the transcriptome-wide JAG1-induced gene expression with that induced by JAG1 + DAPT co-treatment (**Fig. 2C**). By clustering genes according using Pearson correlation, we found that, while two clusters of genes (Cluster A and B) were downregulated by JAG1 treatment, a third cluster (Cluster C) of genes were upregulated in the JAG1 treatment groups and more distinctly in the JAG1 + DAPT-treated group. Because our prior research demonstrated that CNC ossification occurs via a non-canonical NOTCH pathway (8) when treated either with JAG1 or JAG1 + DAPT, these data suggest that Cluster C contains genes involved in pro-ossification pathways.

### JAG1 Canonical and Non-Canonical Induction of Genes Involved in Bone Homeostasis

To determine the individual genes that are both upregulated in JAG1 and JAG1 + DAPT (**Fig. 2C**, Cluster C) and those that were specific to non-canonical JAG1-NOTCH signaling (**Fig. 2C**), we next analyzed the data using volcano plots (**Fig. 3A-C**). Comparing JAG1 to Fc, we found up-regulation of multiple genes within the NOTCH pathway (*Hes1*), bone chemokines (*Cxcl1*), proinflammatory cytokines (*Il6* and *Cxcl12*), cytoskeletal organization regulators (*Rhou*) and, promoter of early osteoblast cell proliferation and inhibitor of osteoclasts (*Id1*) compared to control-treated CNC cells (**Fig. 3D**). To identify genes upregulated by non-canonical JAG1-NOTCH signaling, we next compared JAG1 + DAPT to FC. We found upregulation of *Cxcl1* and *Prl2c2*, (**Fig. 3B & 3D**), which are both involved in bone homeostasis (43, 44). Genes that were upregulated in both JAG1 and JAG1 + DAPT-treated samples, compared to Fc-treated samples, were subsequently compared (**Fig. 3C**). As expected, we also found that CNC cells stimulated with JAG1 in the presence of DAPT demonstrated decreased expression of the classic canonical pathway target, *Hes1* and increased levels of *Prl2c2* (**Fig 3C & 3D**), compared to CNC cells stimulated by JAG1 alone. *Prl2c2*, (encodes Proliferin or Plf1), is a gene known to be upregulated during NOTCH1 haploinsufficiency and is an inducer of cell proliferation (43, 45). *Cxcl1* is a bone chemokine produced by an osteoblast cell to attract osteoclasts to the bone environment in order to trigger bone remodeling (44). *Cxcl1* along with the proinflammatory cytokines, *Il6* and *Cxcl12* was significantly decreased in the presence of DAPT (**Fig. 3C & 3D**) suggesting that JAG1 induces these genes via the NOTCH canonical pathway, while, *Rhou* and *Id1* were preserved in the presence of DAPT suggesting that these genes are induced by the non-canonical JAG1-NOTCH pathway (**Fig. 3C & 3D**).

**Figure 3:**
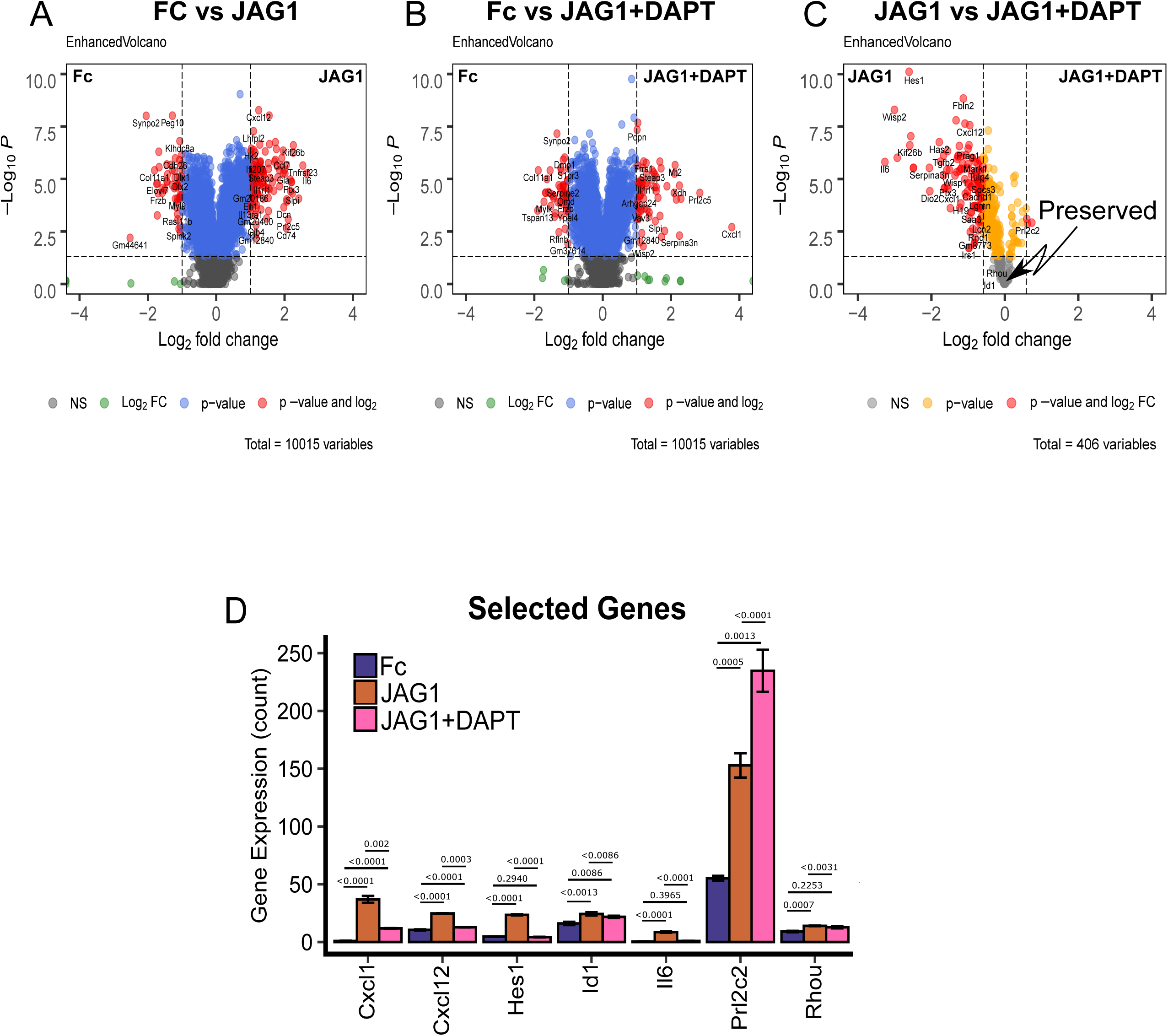
JAG1 induces bone-inductive genes. (A-B) Volcano plots of differentially expressed genes in CNC cells that were treated with JAG1-dynabeads complex (5.7 μM) with or without DAPT (15 µM) compared to Fc-dynabeads complex (5.7 µM). (C) Subsequent comparison of the JAG1 and JAG1 + DAPT treatment groups for genes that were significantly upregulated in either group compared to Fc. Plots indicate genes that were down-regulated in response to DAPT (red) or preserved between groups (grey). (D) Comparison between JAG1 treated-samples in the presence and absence of DAPT revealed various distinct genes, like *Cxcl1, Cxcl12, Hes1* and *Il6* upregulated downstream of JAG1 alone and *Prl2c2* upregulated downstream of JAG1 + DAPT along with some conserved genes like *Id1* and *Rhou*. Data were subjected to ANOVA and Tukey’s post-test and are presented as mean (n ≥ 2) ± SD with p values reported.

### NOTCH Canonical Pathways Are Induced by JAG1 in CNC Cells

To identify gene pathway changes downstream of both canonical and non-canonical NOTCH signaling we utilized the C2 curated gene set annotations from the molecular signatures database (MSigDB) (39) together with Gene Set Variation Analysis (GSVA) (38) to quantify enrichment of 2,199 gene sets among our treatment conditions. Clustering the GSVA values across all gene sets using a Euclidean distance revealed clusters of gene sets that were downregulated by JAG1 + DAPT (Cluster II) and JAG1 alone treatment (Cluster III) and, those that were upregulated (Cluster I) in both JAG1 and JAG1 + DAPT conditions (**Fig. 4A**). The first cluster represents gene sets that are downstream of non-canonical JAG1-NOTCH signaling, while the second represents gene sets downstream of canonical JAG1-NOTCH signaling, that are downregulated non-canonically. These findings indicate that JAG1 stimulates diverse pathways downstream of both canonical and non-canonical notch signaling.

**Figure 4:**
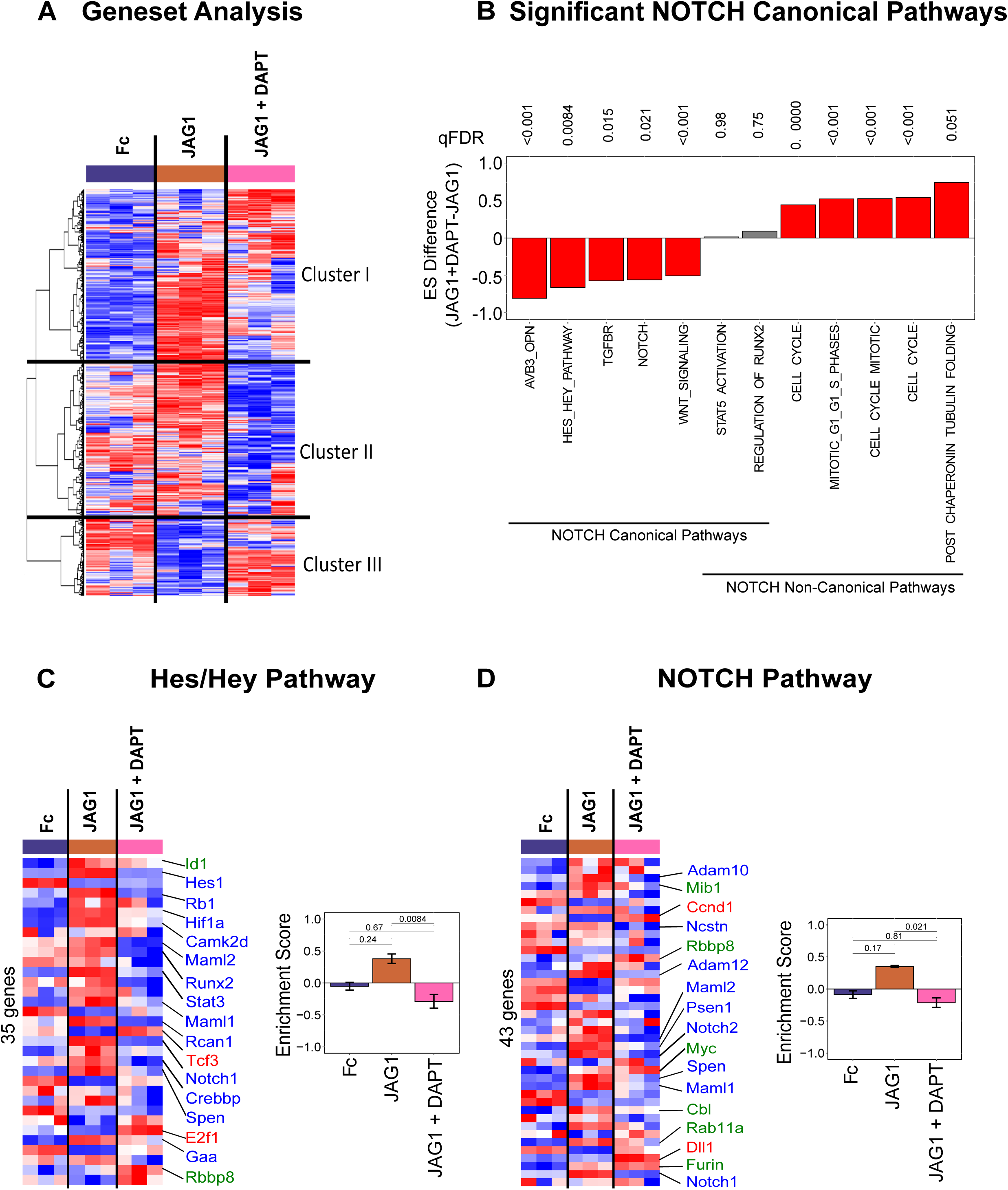
JAG1 enriches canonical NOTCH gene sets. (A) Gene set variation analysis (GSEA) quantified differences in 2199 gene sets. Clustering analysis (Euclidean + Ward’s minimum variance) revealed 3 clusters: Cluster I involves gene sets that were elevated in the JAG1-treated group, Cluster II shows gene sets that were downregulated in JAG1 + DAPT-treated samples and Cluster III indicates gene sets downregulated in JAG1 alone treated cells. (B) GSVA pathway analysis of selected pathways from CNC cells that were treated with JAG1-dynabeads complex (5.7 μM), with or without DAPT and Fc-dynabeads complex (5.7 µM) revealed various pathways up- and downregulated by JAG1 compared to JAG1 + DAPT. The *Hes/Hey* pathway (C) and the NOTCH pathway (D) were upregulated canonically and downregulated non-canonically downstream of JAG1. qFDR values are reported from a permutation analysis. Heatmap labels indicate selected genes whose expression levels are significantly different (p < 0.05) (red, blue, green fonts) in JAG1-treated groups compared to the Fc-treated sample. Expression levels of selected genes are significantly upregulated (Red font), downregulated (Blue font) or unchanged (Green font) in JAG1 + DAPT treated cells compared to JAG1 treated cells. See **Suppl. Fig 1 & 2** for entire list of genes shown in heatmaps from C & D.

Since *Cxcl1, Cxcl12, Hes1, Id1, Il6, Prl2c2* and *Rhou* were significantly upregulated in response to treatment with JAG1 (**Fig. 3D**), we next wanted to determine if any of these previously annotated pathways were associated with canonical NOTCH signaling. We identified upregulation of five NOTCH canonical pathways in JAG1 treatment condition that were significantly downregulated when treated with JAG1 + DAPT (**Fig. 4B**). JAG1 stimulated both the *Hes/Hey* and NOTCH pathways, and these were inhibited in the presence of DAPT, as expected (**Fig. 4C & 4D, Suppl. Table 1 & 2**). Importantly, JAG1 in the presence of DAPT, activated other pathways, such as, the cell cycle, tubulin and the regulators of *Runx2* pathways and downregulated the TGFBR pathway (**Fig. 4B, Suppl. Table 3-5**), a classic BMP2-activated inflammatory and chondrogenic pathway, indicating JAG1’s role in activating bone-inductive pathways without activating the inflammatory pathway via non-canonical NOTCH signaling. Together, these findings affirm that JAG1 stimulates activation of canonical NOTCH pathways, which are inhibited by DAPT.

### NOTCH Non-Canonical Signaling Stimulation by JAG1 Induces Bone Developmental Pathways in CNC cells

Non-canonical NOTCH pathways that were enriched in response to JAG1 or JAG1 + DAPT treatments were examined and we found that JAG1 + DAPT induces various cell-cycle associated pathways, including cyclin-dependent cell cycle genes, *Cdk1, Ccnd1, Cdc25a, Ccne1* and *Cdk6* (**Fig. 4B, 5A, Suppl. Table 3**), indicating that JAG1 promotes CNC cell proliferation via a non-canonical NOTCH pathway. Further, JAG1 + DAPT treatment activated the tubulin pathway, including, *Tuba1c*, and *Tuba4a*, all of which crucially regulate cell structure and membrane reorganization associated with osteoblast commitment (**Fig. 5B, Suppl. Table 4**) (46-48). JAG1 also stimulated expression of regulators of *Runx2* initiation such as, *Smurf1* and *Esrra*, canonically and non-canonically (**Fig. 5C, Suppl. Table 5)**.

**Figure 5:**
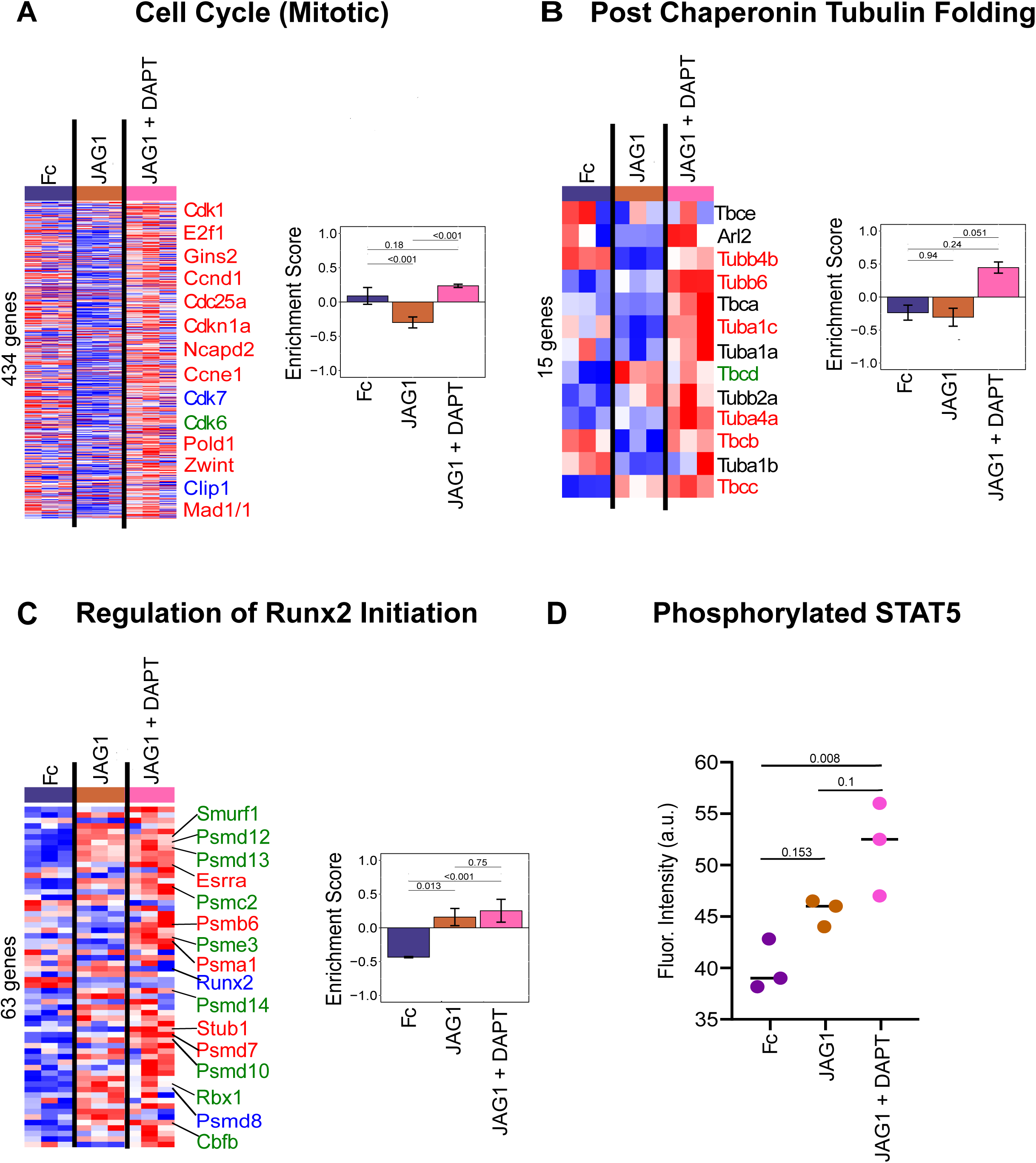
JAG1 enriches bone development gene sets via the NOTCH non-canonical pathway. Pathway analysis of data obtained from sequencing of RNA that was isolated from CNC cells that were treated with JAG1-dynabeads complex (5.7 μM), with or without DAPT, a NOTCH canonical pathway inhibitor and Fc-dynabeads complex (5.7 µM) revealed that JAG1 induces (A) the cell cycle pathway, supporting proliferation, the tubulin pathway, involved in cell structure reorganization essential for CNC cell osteoblast commitment, the regulation of *Runx2* initiation pathway, essential for osteoblast proliferation and commitment. Heatmap labels indicate selected genes whose expression levels are significantly different (p < 0.05) (red, blue, green fonts) in JAG1-treated groups compared to the Fc-treated sample. Those in black font are not significantly different from Fc-treated groups (p > 0.05). Expression levels of selected genes are significantly upregulated (Red font), downregulated (Blue font) or unchanged (Green font) in JAG1 + DAPT treated cells compared to JAG1 treated cells. See **Suppl. Fig 3, 4 & 5** for entire list of genes. qFDR values are reported from a permutation analysis. (D) Luminex analysis revealed that STAT5 phosphorylation was increased by JAG1 treatment. Data were subjected to ANOVA and Tukey’s post-test and are presented as mean (n ≥ 2) ± SD with p values reported.

STAT5 is a known bone regulatory pathway downstream of JAK2 signaling (49) and thus, we tested its activation in CNC cells treated with JAG1 + DAPT. A Luminex assay (Millipore) was used to quantify STAT5 phosphorylation in CNC cells treated with JAG1 ± DAPT, revealing increased phospho-STAT5 in the JAG1 treatment condition with DAPT (**Fig. 5D**). Together these data show that JAG1 up-regulates cell proliferation, ossification and signaling pathways responsible for bone formation and regeneration through non-canonical JAG1-JAK2-STAT5 signaling (**Fig. 6**).

**Figure 6:**
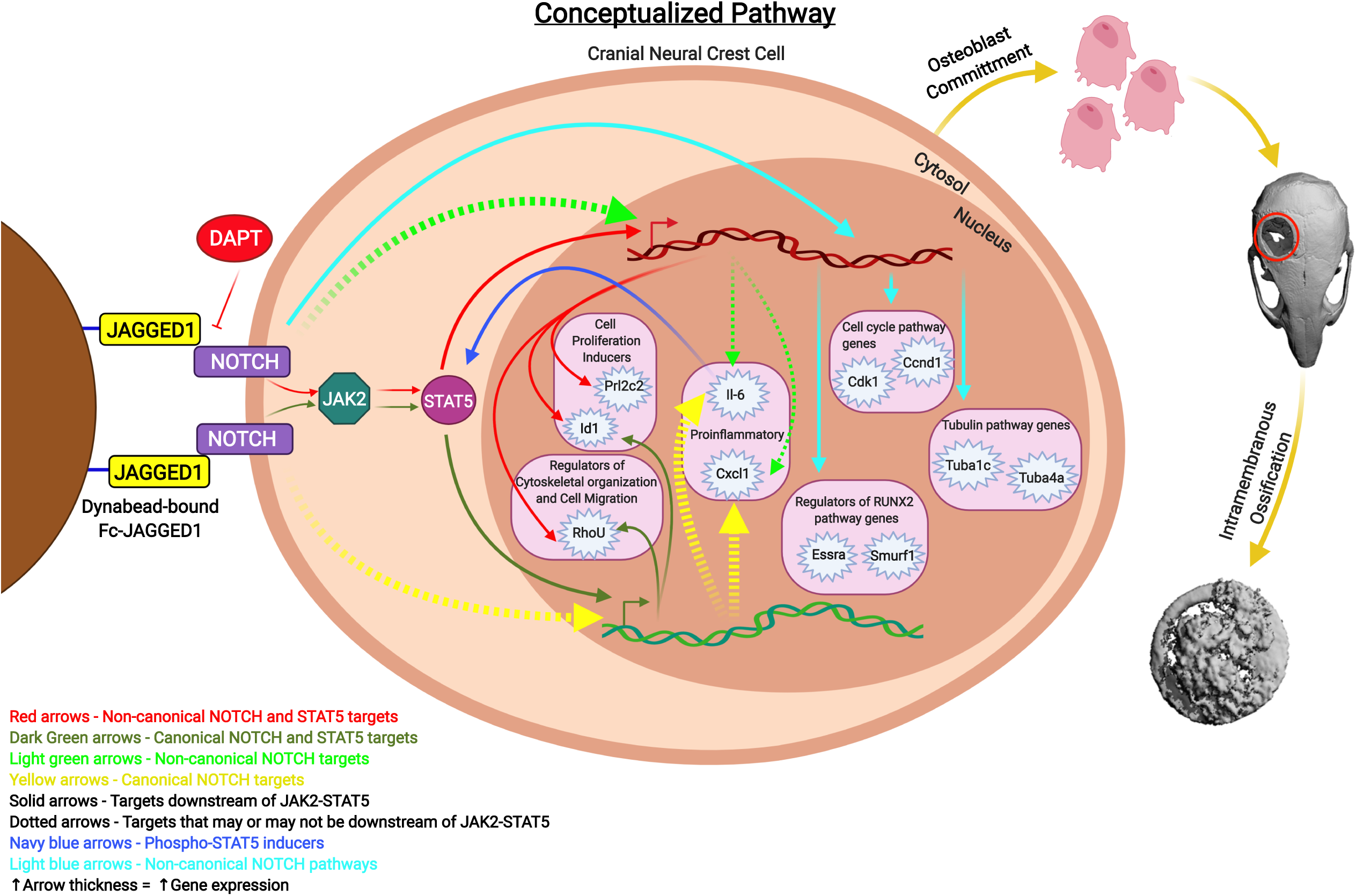
Summary & conceptualized pathway. Cranial neural crest cells treated with JAG1-dynabeads with or without DAPT, a NOTCH canonical pathway inhibitor, showed activation of the JAK2-STAT5 pathway. Red and Green solid arrows indicate non-canonical and canonical NOTCH targets of JAG1, respectively, downstream of JAK2-STAT5 (*Prl2c2, Id1, Rhou*). Light green arrows indicate non-canonical NOTCH targets and yellow arrows indicate canonical NOTCH targets. Dotted arrows indicate targets that may or may not be activated by STAT5. Navy blue arrows indicate targets that can induce phosphorylation of STAT5. Pathways activated non-canonically downstream of JAG1 revealed by pathway analyses are indicated by light blue arrows. Greater the arrow thickness higher the gene expression levels. All these events lead to JAG1-induced CNC cell osteoblast commitment and thereafter bone formation by intramembranous ossification.

## 4. Discussion

To our knowledge, this is the first study to holistically define the activation of the non-canonical NOTCH pathway in response to JAG1 and explore its potential for bone regeneration. JAG1 is required for multiple developmental processes, including normal craniofacial development, based on its role in cell fate determination (50). NOTCH signaling occurs in a cell-to-cell manner where a membrane bound protein ligand (JAG1, JAG2, DLL1-4) binds to the NOTCH receptors 1-4 on an adjacent cell (51). Mutations in *JAG1* signaling are associated with Alagille Syndrome in humans and lead to cardiac, biliary, bony and craniofacial abnormalities (52). Prior research performed by our lab and others have investigated the role of JAG1 in craniofacial development and found that absence of *Jag1* in mice leads to craniofacial bone loss, a phenocopy of Alagille Syndrome patients (4, 13, 53).

There are currently no available craniofacial bone regenerative approaches for children. Our goal was to characterize the osteo-inductive properties of JAG1 and harness its potential as a bone regenerative therapy. As a proof-of-concept, we formulated a hydrogel encapsulating CNC cells to deliver JAG1 to regenerate bone in critical-sized calvarial defects in juvenile mice. Hydrogels are an established delivery system due to their diverse compositions and adjustable physicochemical properties (54, 55). In the JAG1-PEG-4MAL hydrogel, JAG1 was immobilized onto dynabeads coated with protein G to recreate the appropriate JAG1 orientation for cell-to-cell NOTCH signaling to occur. Delivery of the JAG1-PEG-4MAL-CNC hydrogel significantly improved closure of the critical-sized defect at 20 µM of JAG1 and 10 µM had significant bone deposition as well (**Fig. 1I**). There are previous reports of bone induction using JAG1 delivery *in vivo* using a collagen sponge, similar to the delivery of BMP2 in humans (5), however it is was not determined if JAG1 was bound in an orientation to facilitate cell-to-cell signaling (5). In our study, delivery of BMP2 alone demonstrated increased ossification in cranial critical-sized defects (**Suppl Fig. 1**). However, BMP2 with CNC cells showed minimal bone regeneration, suggesting that encapsulation of the BMP2 with CNC cells may have degraded or inhibited the BMP2. Also, CNC cells have been found to need BMP2 prior to migration from the dorsal tube, but once they reach the branchial arches and are prepared for osteoblast commitment, the CNC cells may be less responsive to BMP2 (42).

The development of the JAG1-PEG-4MAL-CNC hydrogel provides a proof-of-concept bone regenerative strategy. However, this method cannot be extrapolated to humans since dynabeads are super-magnetized particles that could have off-target effects. Future studies will focus on the direct conjugation of JAG1 to the PEG-4MAL hydrogel as a step towards a regenerative therapy for children with craniofacial bone defects. The exact mechanism by which JAG1 induced osteoblast commitment in the CNC cells was unknown, and we used RNAseq to determine the downstream targets of non-canonical JAG1 signaling.

We previously showed that JAG1 non-canonically signals through NOTCH via phosphorylation of JAK2 during CNC cell osteoblast commitment (8). In this manuscript, we demonstrated increased STAT5 phosphorylation, demonstrating that STAT5 (**Fig. 5D**), a known bone inductive phospho-protein (56), is a downstream target of the JAG1-JAK2 axis during pathway inhibition in the presence of DAPT. STAT5 phosphorylation is involved in osteoblast commitment and it is upstream of multiple osteoblast regulatory genes including *Runx2, Igf1, Bmp2* and *Tbx3* (56). We found that JAG1 treatment with DAPT repressed canonical NOTCH targets *Hes1* and *Hey1* and increased expression levels of *Prl2c2*, which are typically repressed by canonical NOTCH signaling, and is a potent inducer of cellular proliferation (**Fig. 3D & 4C**) (57). *Prl2c2* is also a downstream target of phospho-STAT5, suggesting that increased phosphorylation of STAT5 is responsible for increased *Prl2c2* expression (58). These data suggest that the non-canonical JAG1-JAK2-STAT5-*Prl2c2* signaling axis is responsible for CNC cell proliferation.

Increased expression of *Id1* in the CNC cells following JAG1 delivery was observed despite the presence of DAPT, suggesting that it is induced through the non-canonical pathway (**Fig. 3C & 3D**). Id1 is a helix-loop-helix protein that is involved in the early osteoblast proliferation and cell fate commitment (59). In erythroid cells, JAK2-STAT5 signaling plays a key role during normal hematopoiesis and STAT5 was found to bind and transactivate a downstream enhancer of *Id1* (60), suggesting that JAG1-JAK2-STAT5-*Id1* is similarly an active pathway in the CNC cells. An additional STAT5-*Id1* pathway function involves the inhibition of osteoclast formation which indirectly adds to bone formation, and that knockout of the STAT5 pathway is associated with osteoporosis (61).

GTPase RhoU (*Rhou*) is known to be involved in mediating cytoskeletal dynamics and has also been shown to be a downstream target of the JAK-STAT pathway in myeloma cells where it facilitates cell proliferation and migration (62). *Rhou*, similarly to *Id1*, reduces osteoclast differentiation (63). *Rhou* expression remains increased when CNC cells are treated with JAG1 with or without DAPT (**Fig. 3D**), suggesting that JAG1 regulates the migration and cytoskeletal changes of CNC cells during osteoblast commitment through the JAG1-JAK2-STAT5-*Rhou* non-canonical pathway.

JAG1 induced CNC cells show a robust increase in expression of *Cxcl1* and *Il6* (**Fig. 3C & 3D**) compared to that in CNC cells treated with JAG1 in the presence of DAPT. CXCL1 is a bone chemokine produced by an osteoblast cells to attract osteoclasts to the bone environment in order to trigger bone remodeling (44). IL-6, is a proinflammatory cytokine that induces T-cell migration and negatively regulates osteoblasts, both leading to reduced bone formation (64, 65). Thus, *Cxcl1* and *Il6*, which induce bone resorption and inflammation, respectively, are canonical NOTCH pathway targets which suggest that harnessing the non-canonical JAG1 signals can avoid the osteolytic and osteoinhibitory pathways downstream of canonical NOTCH signaling.

Commitment of CNC cells to osteoblasts is accompanied by changes in cell structure where the cytoplasm increases 26-fold with the increased production of intracellular filaments and microtubules (66). Pathway analysis following JAG1 + DAPT treatment of CNC cells (**Fig. 5B**) demonstrated significant changes in the tubulin pathways genes, *Tuba1c, Tuba4a*, (46-48) suggesting that multiple cytoskeletal genes are downstream of JAG1-JAK2-STAT5 signaling. Since prior studies have demonstrated the importance of cytoskeletal changes during osteoblast development and specifically the presence of microtubules within cellular processes and the transition of the processes as osteoblasts become osteocytes (67) our future studies will characterize the various cytoskeletal filaments and microtubules that change in response to JAG1 treatment.

The first step in osteoblast formation requires a quiescent pre-osteoblast cell to enter the cell cycle and undergo proliferation. Following this proliferative stage, expression of *Runx2* promotes osteoblast lineage commitment during which it encourages the pre-osteoblasts to exit from the cell cycle (68). In our prior work we found that osteogenic cultures of CNC cells treated with JAG1 + DAPT have an initial proliferative phase for the first week and then express *Runx2* on day 10 to allow CNC cell osteoblast commitment (8). In this paper, our RNAseq data after 12 hours of JAG1 + DAPT treatment of CNC cells revealed that cell cycle genes, such as *Cdk1, Ccnd1, Cdc25a, Ccne1* and *Cdk6* were upregulated, indicating that JAG1 + DAPT treatment had induced cell proliferation (69, 70). Moreover, we observed an upregulation of *Smurf1* and *Esrra* along with a downregulation of *Runx2*. (**Fig 5A & C**). This observation is consistent with other studies that report a transient inhibition of *Runx2* expression by *Smurf1*, which allows the preosteoblastic cell to remain in the cell cycle to undergo proliferation (71). These results suggest that CNC cells are induced in preparation for osteoblast commitment with the anticipated expression of osteoblast genes, like *Runx2*, in the following week. Other researchers have also noted that increased expression of *Esrra* leads to increased *Runx2* expression and induces osteoblast differentiation, suggesting that timing of the proliferative phase prior to the osteoblast commitment phase is critical (72, 73).

Altogether, our data demonstrate that a proof of principle JAG1-PEG-4MAL-CNC hydrogel regenerates bone reliably in a cranial defect model. We have also identified an important non-canonical NOTCH JAG1-JAK2-STAT5 pathway (**Fig. 6**) that controls multiple cell cycle genes (*Cdk1, Ccnd1, Cdc25a, Ccne1* and *Cdk6)* and those that promote preosteoblasts to enter the proliferative phase (*Prl2c2, Id1* and *Smurf1*), to undergo cytoskeletal changes (*Tuba1c, Tuba4a* and *Rhou*) and to commit to the osteoblast lineage (*Esrra*). Future studies will focus on elucidating the additional downstream pathway targets of this non-canonical system and developing novel bone regenerative therapies based on this information. Moreover, understanding the intracellular cytoskeletal changes during JAG1-induced CNC cell osteoblast commitment will expand on the role of non-canonical NOTCH signaling on cell morphology and differentiation.

## Abbreviations

*Ccnd1*: Cyclin D1
*Cdc25a*: Cell Division Cycle 25a
*Cdk1*: Cyclin Dependent Kinase 1
CNC: Cranial Neural Crest
*Cxcl1*: Chemokine C-X-C Motif Ligand 1
*Esrra*: Estrogen Related Receptor Alpha
*Hes1*: Hairy and Enhancer of Split 1
*Id1*: Inhibitor of DNA Binding 1
*Il6*: Interleukin 6
JAG1: JAGGED1
JAK2: Janus Kinase 2
*Prl2c2*: Proliferin
*Rhou*: Ras Homolog Family Member U
RNAseq: Ribonucleic Acid Sequencing
*Runx2*: Runt-related Transcription Factor 2
*Smurf1*: Smad Ubiquitin Regulatory Factor 1
STAT5: Signal Transducer and Activator of Transcription 5
*Tuba4a*: Tubulin Alpha 4a

## 5. Acknowledgements

Study design: AK, JMM, NJW, HMD, AJG, LBW and SLG. Study conduct: AK. Data collection: AK, JMM, AMA, SAB and PB. Data analysis: AK, JMM, DSD, AMA, SAB, PB and LBW. Data interpretation: All authors. Drafting manuscript: AK, JMM, LBW and SLG. Revising manuscript content: All authors. Approving final version of manuscript: All authors. AK, LBW and SLG take responsibility for the integrity of the data analysis. Research reported in this publication was supported by the Oral Maxillofacial Surgery Foundation (Funding ID: 2591), and National institute of Health, National Institute of Arthritis and Musculoskeletal and Skin Diseases of the National Institutes of Health under Award Numbers DE026762, R01AR062920 and R01AR062368, respectively. This manuscript has not been copyrighted or published before and is not under consideration for publication elsewhere. We also assure you that the contents of this manuscript will not be subsequently published in the same or similar form in any language without your consent. Lastly, the authors declare no conflict of interest.

## 7. Figure Legends

**Supplemental Figure 1:**
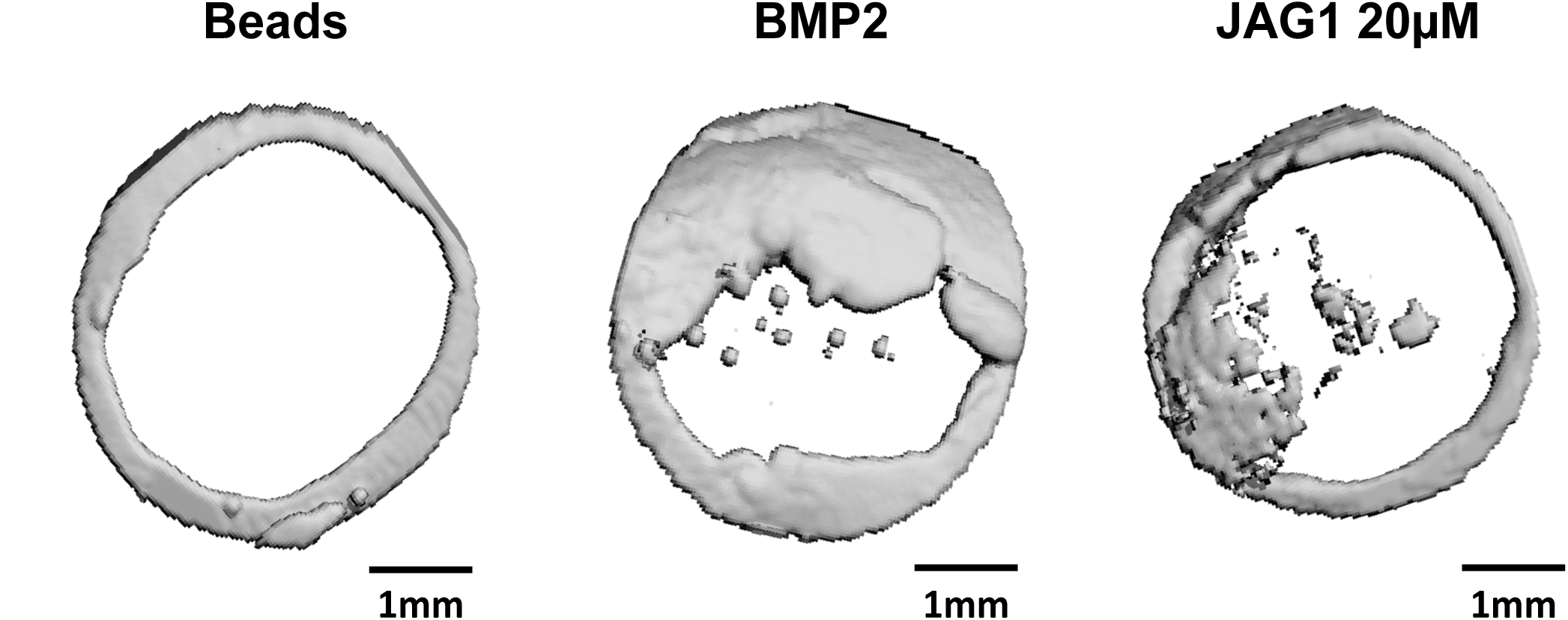
μCT 3D reconstructions, 12 weeks after 3.5 mm critical-sized defects were created in mouse parietal bone and implanted with 4% PEG-MAL hydrogels incorporated with JAG1-dynabeads complex (5 μM, 10 μM or 20 μM), dynabeads alone and BMP2 (2.5 µM) as 3 separate doses (Initial dose, Week 4, Week 8).

**Supplemental Table 1:** List of genes involved in the *Hes1/Hey1* pathway along with expression differences between Fc-, JAG1- and JAG1 + DAPT-treated samples. Significant difference is indicated by p < 0.05.

**Supplemental Table 2:** List of genes involved in the NOTCH pathway along with expression differences between Fc-, JAG1- and JAG1 + DAPT-treated samples. Significant difference is indicated by p < 0.05.

**Supplemental Table 3:** List of genes involved in the cell cycle pathway along with expression differences between Fc-, JAG1- and JAG1 + DAPT-treated samples. Significant difference is indicated by p < 0.05.

**Supplemental Table 4:** List of genes involved in the post chaperonin tubulin folding pathway along with expression differences between Fc-, JAG1- and JAG1 + DAPT-treated samples. Significant difference is indicated by p < 0.05.

**Supplemental Table 5:** List of genes involved in the regulation of *Runx2* pathway along with expression differences between Fc-, JAG1- and JAG1 + DAPT-treated samples. Significant difference is indicated by p < 0.05.

